# Common features in plastic changes rather than constructed structures in recurrent neural network prefrontal cortex models

**DOI:** 10.1101/181297

**Authors:** Satoshi Kuroki, Takuya Isomura

**Affiliations:** Laboratory for Behavioral Genetics, RIKEN Brain Science Institute, Wako, Saitama 351-0198, Japan; Laboratory for Neural Computation and Adaptation, RIKEN Brain Science Institute, Wako, Saitama 351-0198, Japan

## Abstract

We have flexible control over our cognition depending on the context or surrounding environments. The prefrontal cortex (PFC) controls this cognitive flexibility; however, the detailed underlying mechanisms remain unclear. Recent developments in machine learning techniques have allowed simple recurrent neural network PFC models to perform human- or animal-like behavioral tasks. These systems allow us to acquire parameters, which we could not in biological experiments, for performing the tasks. We compared four models, in which a flexible cognition task, called context-dependent integration task, was performed; subsequently, we searched for common features. In all the models, we observed that high plastic synapses were concentrated in the small neuronal population and the more concentrated neuronal units contributed further to the performance. However, there were no common properties in the constructed structures. These results suggest that plastic changes can be more general and important to accomplish cognitive tasks than features of the constructed structures.

## Introduction

Humans can generate and flexibly switch cognitive sets depending on the situation even with the same sensory-sensory or sensory-motor associations. The prefrontal cortex (PFC) controls cognitive flexibility, also the called executive function (Miller & Cohen, 2001; Nobre & Kastner, 2014). Although the control mechanism of executive function has been studied using animal models, the detailed underlying mechanism remains unclear. One of the reasons is the limited number of observable and manipulatable biological variables of the brain. Recent developments in simple recurrent neural network (RNN) computational models allowed the study of behavioral tasks and measurement of cognitive flexibility in animal models. The network activities resembled those the PFC activities during the performance of similar behavioral tasks (Mante, Sussillo, Shenoy, & Newsome, 2013; Miconi, 2017; Song, Yang, & Wang, 2016, 2017). With this novel computational method, we are now able to analyze nearly all the parameters involved in behavioral tasks.

Mante et al. developed a behavioral task for evaluating the cognitive flexibility of monkeys by modifying a random-dot motion task and adding color factors to the motion factors (Mante et al., 2013), called context-dependent integration task (Song et al., 2016). For the task, the monkey needed to select one of two choices depending on colored dots moving randomly on a screen. The contextual information of the task was changed regarding whether the monkey should select a correct answer based on color or motion (Figure 1A). In addition, Mante et al. constructed an RNN model for performing the task, in which populational activities during the task were similar to those of the monkey’s PFC neurons. However, the optimization method used for the model was hessian-free (HF) optimization (HF model) (Martens, 2010; Martens & Sutskever, 2011), which is not sufficiently biologically plausible.

**Figure 1:**
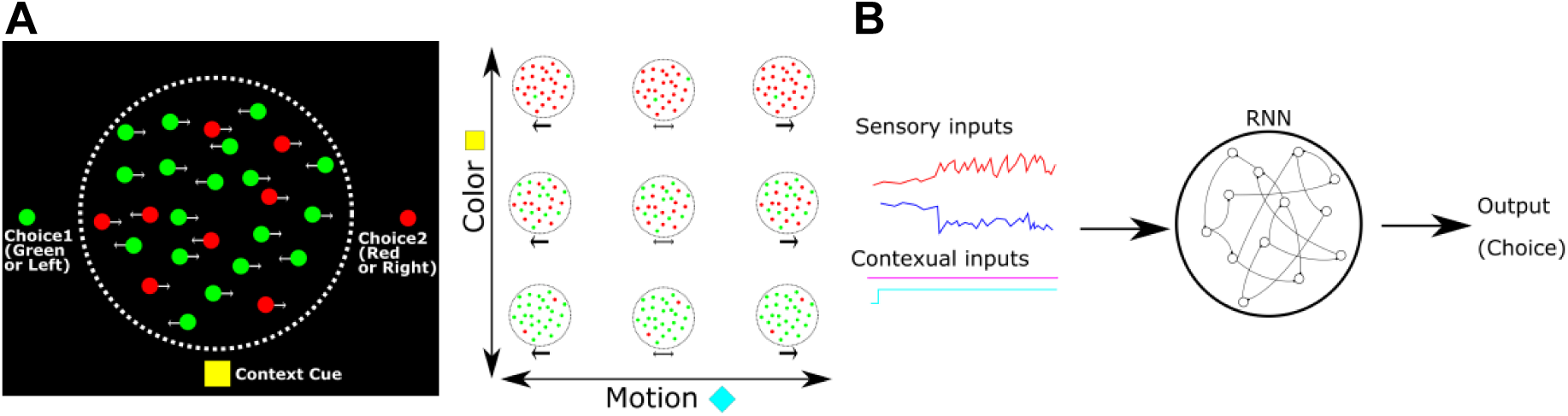
(A) Context-dependent integration task applied to a monkey. The left panel shows a schematic of the task. At the center of the screen, the green or red dots move randomly from right to left. The monkey should choose between the left green or right red option by saccade while referring to the central moving dots. The reference (color or motion) for the correct answer depended on the contextual cue at the bottom of the screen. The right panel indicates a variety of colored moving dot patterns and contextual cues.(B) Schematic image of RNN PFC models. All models consist of input sensory and contextual information and output choices. In the RNN, neuronal units are connected to each other.

Aiming for more biologically plausible models, a few groups suggested other RNN models for performing context-dependent integration tasks. Song et al. proposed an RNN model (pycog model) (Song et al., 2016), which comprised separate excitatory and inhibitory neuronal units and involved the use of a simpler optimization method than HF, namely, a modified stochastic gradient descent (SGD) (Pascanu, Mikolov, & Bengio, 2013). In addition to context-dependent integration tasks, the model allowed the study of several PFC dependent behavioral tasks.

Song et al. constructed another RNN model (pyrl model), which consisted ofa policy (choosing next behaviors) and a baseline network (evaluating feature rewards) by which learning was reinforced with reward signals and not error signals between actual and optimal outputs (Song et al., 2017). This method was built with policy gradient reinforcement learning, particularly the REINFORCE algorithm for RNN (Wierstra, Forster, Peters, & Schmidhuber, 2010; Williams, 1992), and the structure was known as an actor-critic structure (Sutton & Barto, 1998). The system optimized the baseline network to estimate the future rewards in each situation and the policy network to make an optimal choice in order to maximize the future rewards. This model consisted of value-evaluating tasks as well as cognitive tasks.

Miconi introduced the reward-modulated Hebbian rule as an optimization method (rHebb model) (Miconi, 2017). He introduced a reward-modulated Hebbian rule and node-perturbation method for learning (Fiete, Fee, & Seung, 2007), assuming that the algorithm is more biologically plausible than HF or SGD. The system also allowed the study of several cognitive tasks.

We compared the synaptic weight structures of these four RNN models (HF, pycog, pyrl, and rHebb models) while performing context-dependent integration tasks (Figure 1B) and searched for common features among these models, assuming that the general features could be important to perform the tasks. Although we could not detect any common structures among the learned network structures, we found interesting features in plastic change from the initial network state to the last learned state, which was conserved among all models: (1) plastic synapses were concentrated in small populations of units, particularly postsynaptically, meaning that the unital distribution was highly skewed; (2) distributions of the synaptic weight changes had high kurtosis. Additionally, the highly plastic units had greater contributions to performing behaviors than the low plastic units in the HF, pycog, and pyrl but not in the rHebb model. As far as we know, this study is the first to report a detailed structural analysis of the RNN PFC models. The present results indicate that plastic changes secondary to task learning can be more important than the last established structure of the system and can give us new insights on biological and computational neuroscience research.

## Results

### Analysis of the constructed structures

At first, we confirmed whether each system could allow the study of context-dependent integration task (Figure 2A). All systems showed psychometric curves (the relationship between sensory inputs and behavior responses), which changed depending on the context information. More than 85% of the results were correct. Next, the synaptic weight values of each learned system were visualized (Figure 2B). We used excitatory-excitatory (E-E) connections in the pycog model for the analysis because this is the most popular connection in the model. Additionally, we used the policy network in the pyrl model because the baseline network was not related to the choice behavior even though it is important to learn the task (see Materials and Methods). We detected line patterns in the HF model, meaning that high negative or positive weight seemed to be concentrated in few portions of the neuronal units, particularly postsynaptically. However, we could not find any specific patterns in the other models.

**Figure 2:**
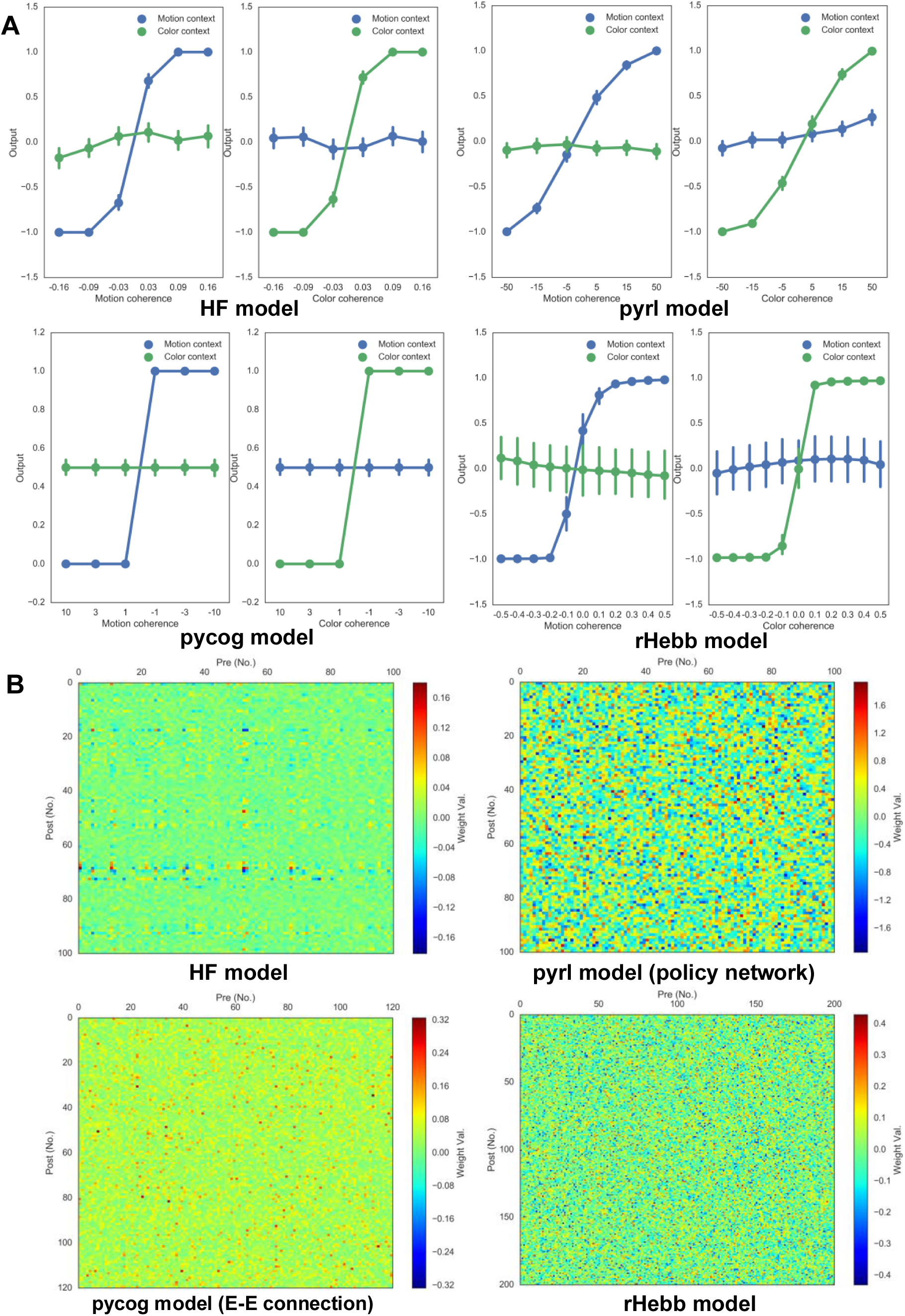
(A) Task performance and (B) synaptic weight values from pre- (horizontal axis) to post-unit (vertical axis) of each system after task learning. We plotted distributions of the synaptic weight values (Figure 3 and Table 1). HF and rHebb had a Gaussian distribution as initial states. The HF system showed non-Gaussian high-kurtosis distribution after task learning, whereas rHebb still showed Gaussian distribution. The pycog and pyrl models did not show Gaussian distributions even from the initial states (see Materials and Methods). The initial distributions are shown in Figure 3-Figure supplement 1.

Because both negative and positive high weights seemed concentrated in sparse unital populations postsynaptically in the HF model, we sorted the units by means of absolute weight values per post-units (defined as post-unital mean weight) and confirmed that high weights were postsynaptically concentrated to a few units in the HF model (Figure 4A). To quantify the weight concentration in the sparse population, we analyzed the distribution of the post-unital mean weight values. The distribution of the HF model showed a highly skewed distribution (Figure 4B and Table 2). The other models did not show that kind of clear weight concentration. In conclusion, with learned network analysis, we could not find any common features in static network structures across all four models, even though all of them succeeded at the learning task.

**Table 1:** Distribution properties in learned network weights

| Model | n | Normality |  | Skewness |  |  | Kurtosis |  |  |
| --- | --- | --- | --- | --- | --- | --- | --- | --- | --- |
|  |  | p | W | p | Z | skew. | p | Z | kurt. |
| HF | 10,000 | 0.00 | 0.93 | 0.00 | 13.9 | 0.35 | 0.0 | 35.9 | 6.17 |
| pycog (E-E) | 14,400 | 0.00 | 0.89 | 0.00 | 53.8 | 1.46 | 0.00 | 33.7 | 3.78 |
| pyrl (policy) | 10,000 | 0.00 | 0.97 | 0.94 | -0.08 | 0.00 | 0.00 | -34.1 | -0.87 |
| rHebb | 40,000 | 0.99 | 0.99 | 0.52 | -0.64 | -0.01 | 0.94 | 0.06 | 0.00 |

**Table 2:** Distribution properties in post-unital mean weight values of the learned network

| Model | n | Normality |  | Skewness |  |  | Kurtosis |  |  |
| --- | --- | --- | --- | --- | --- | --- | --- | --- | --- |
|  |  | p | W | p | Z | skew. | p | Z | kurt. |
| HF | 100 | 0.00 | 0.81 | 0.00 | 5.33 | 1.64 | 0.00 | 3.21 | 2.53 |
| pycog (E-E) | 120 | 0.60 | 0.99 | 0.39 | 0.85 | 0.81 | 0.48 | 0.71 | 0.19 |
| pyrl (policy) | 100 | 0.23 | 0.98 | 0.26 | 1.13 | 0.26 | 0.34 | 0.95 | 0.32 |
| rHebb | 196 | 0.55 | 0.99 | 0.34 | 0.96 | 0.16 | 0.34 | 0.95 | 0.26 |

**Figure 4:**
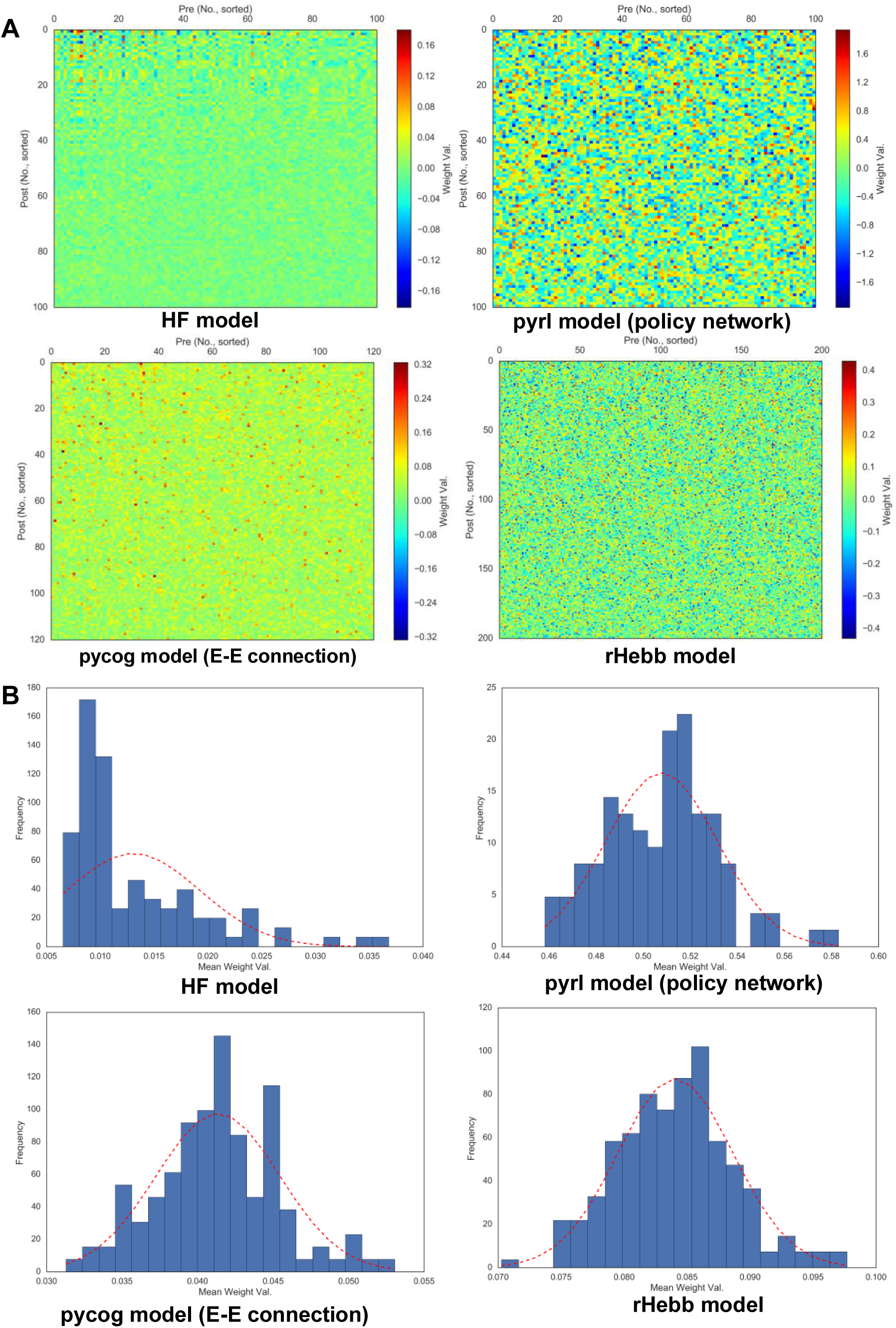
(A) Weight value color plots sorted by post-unital mean weights. (B) Distribution of post-unital mean weights. Doted red line indicates normal distribution with mean and sigma of the values.

### Analysis of plastic changes with task learning

We then analyzed synaptic weight changes by task learning defined as the difference between the initial and final weight values. We found that the weight changes of all models showed line pattern-like HF learned weight distributions as shown in Figure 2B. For a few units, the weight changes tended to be larger either as increases or decreases compared with other units. We sorted the neural units by post-unital mean weight changes (Figure 5A), similar to the post-unital mean weight values of the learned structures (Figure 4A). All the models showed concentrated weight changes in sparse neural populations. By quantitatively analyzing the distribution of post-unital mean weight changes, we found that all models showed highly skewed distributions. This indicates that the networks consisted of a small number of high-plasticity units and a large number of low-plasticity units (Figure 5B and Table 3). Notably, only 10% of the synapses in the pyrl model were plastic (the other synapses did not change through learning) as a default setting.

**Table 3:** Distribution properties in post-unital mean weight changes

| Model | n | Normality |  | Skewness |  |  | Kurtosis |  |  |
| --- | --- | --- | --- | --- | --- | --- | --- | --- | --- |
|  |  | p | W | p | Z | skew. | p | Z | kurt. |
| HF | 100 | 0.00 | 0.87 | 0.00 | 4.44 | 1.25 | 0.04 | 2.08 | 1.16 |
| pycog (E-E) | 120 | 0.00 | 0.93 | 0.00 | 4.09 | 1.01 | 0.05 | 1.97 | 0.97 |
| pyrl (policy) | 100 | 0.00 | 0.91 | 0.00 | 4.54 | 1.29 | 0.00 | 3.06 | 2.31 |
| rHebb | 196 | 0.00 | 0.96 | 0.00 | 4.16 | 0.78 | 0.07 | 1.81 | 0.68 |

**Figure 5:**
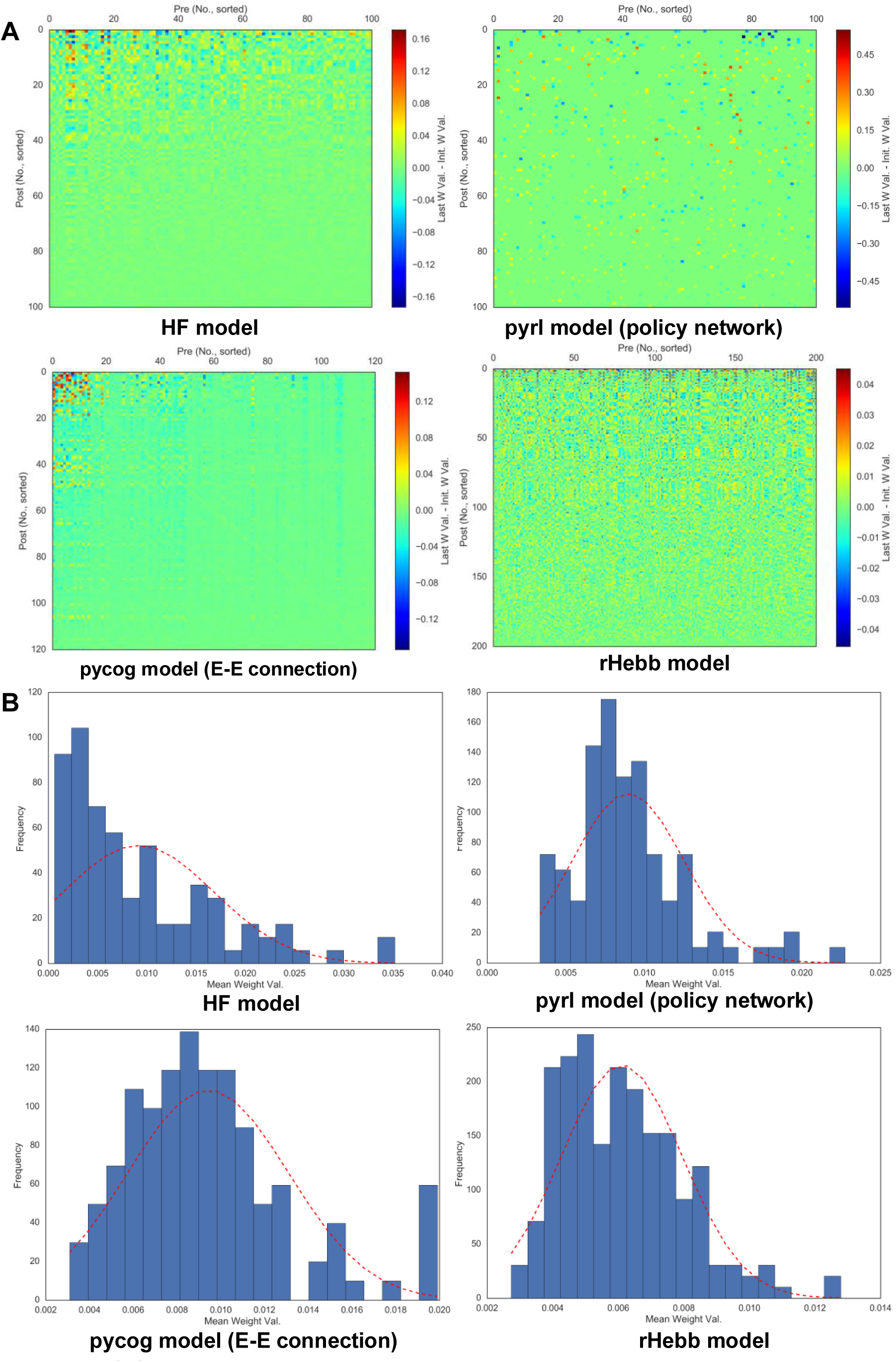
(A) Color plots of weight changes sorted by post-unital mean weight changes. (B) Distribution of post-unital mean weight changes. The dotted red line indicates a normal distribution of the mean and sigma of the values.

In addition, the weight difference distributions of all the models were high-kurtosis distributions as results of the statistical tests, while the shapes of the distributions of pyrl and rHebb model were close to a Gaussian distribution (Figure 6 and Table 4). Pycog and pyrl models had other network connections, such as inhibitory networks in pycog and baseline networks in pyrl. Most of these connections similarly showed that large weight changes were concentrated in the restricted populations of neuronal units (Figure5-Figure supplement 1).

**Table 4.** Distribution properties in weight changes.

| Model | n | Normality |  | Skewness |  |  | Kurtosis |  |  |
| --- | --- | --- | --- | --- | --- | --- | --- | --- | --- |
|  |  | p | W | p | Z | skew. | p | Z | kurt. |
| HF | 10,000 | 0.00 | 0.83 | 0.00 | 22.6 | 0.60 | 0.00 | 43.1 | 11.1 |
| pycog (E-E) | 14,400 | 0.00 | 0.62 | 0.00 | 85.4 | 3.40 | 0.00 | 66.1 | 36.8 |
| pyrl (policy) | 1,000 | 0.00 | 0.99 | 0.00 | -2.82 | -0.22 | 0.00 | 5.30 | 1.24 |
| rHebb | 38,416 | 0.00 | 0.99 | 1.00 | 0.01 | 0.00 | 0.00 | 6.04 | 0.16 |

**Figure 6:**
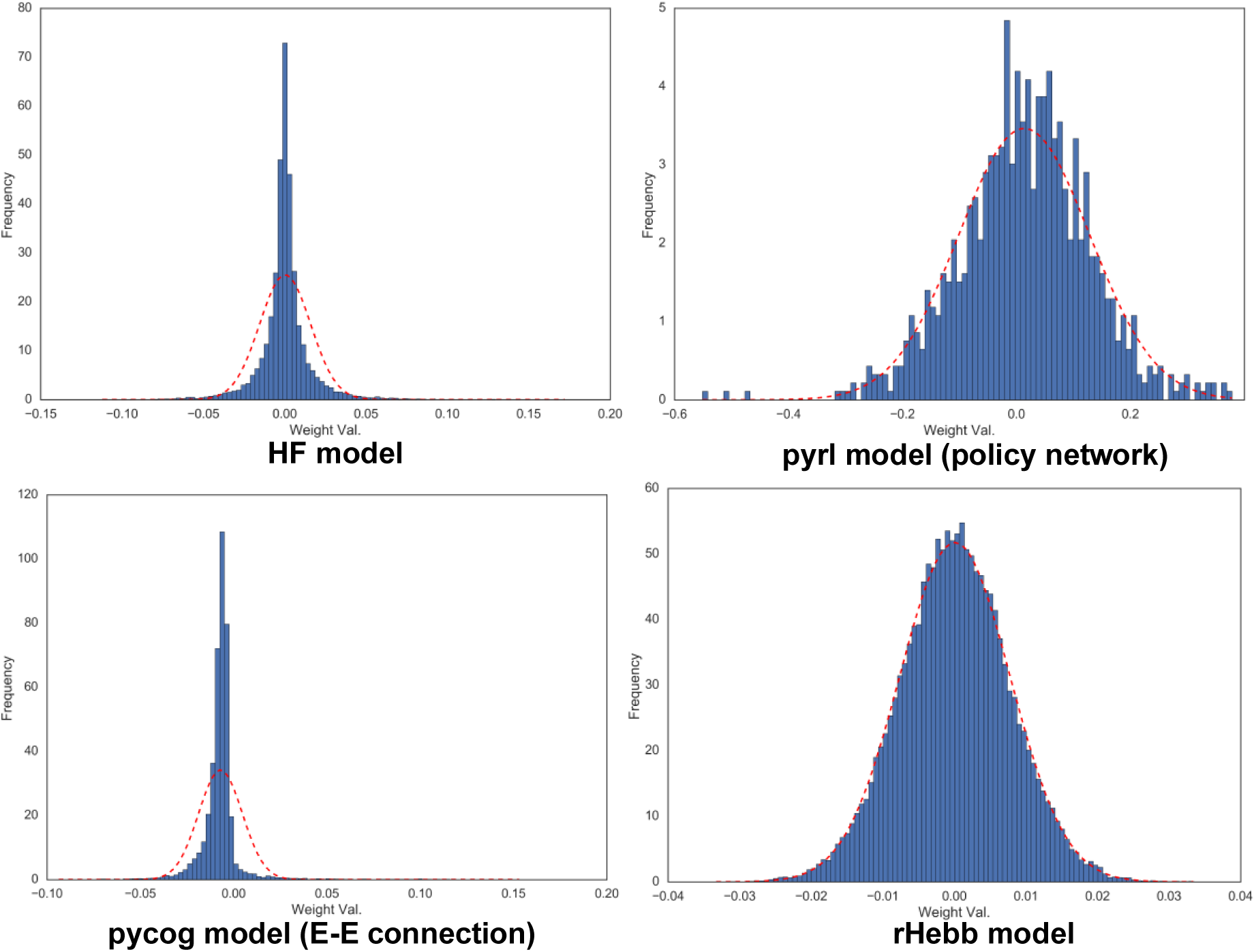
Distribution of weight changes. The dotted red line indicates a normal distribution of the mean and sigma of the values.

We validated whether units with higher plasticity had higher contributions to behavior performances. The fixed number of units in each model was inactivated (n_inact) with ascending (starting from low-plasticity units), descending (starting from high-plasticity units) and shuffled order (sort_type) based on sorted values as shown in Figure 5A while the behavioral task was being performed (Figure 7 and Table 5). The “n” in Table 5 indicates the number of systems used for the test with different initial settings from different random seeds. All the models showed significant differences in behavior performances among sort_type x n_inact interaction and/or sort_type in a two-way analysis of variance (ANOVA), but there was no significant difference among sort type in rHebb model with post-hoc multiple comparisons. These results indicated that units with higher plasticity in HF, pycog, and pyrl models made larger contributions to task performance; however, the rHebb model presented redundancy for the loss of the high-plasticity units.

**Table 5:** Two-way ANOVA results of the inactivation experiments

| Model | test | df (factor) | df (error) | F | p |
| --- | --- | --- | --- | --- | --- |
| HF<br>(n = 11) | n_inact | 9 | 300 | 34.9 | 0.00 |
|  | sort_type | 2 | 300 | 231 | 0.00 |
|  | n_inact × sort_type | 18 | 300 | 4.87 | 0.00 |
| pycog<br>(n = 11) | n_inact | 11 | 360 | 50.6 | 0.00 |
|  | sort_type | 2 | 360 | 355 | 0.00 |
|  | n_inact × sort_type | 22 | 360 | 9.63 | 0.00 |
| pyrl<br>(n = 20) | n_inact | 9 | 570 | 596 | 0.00 |
|  | sort_type | 2 | 570 | 24.6 | 0.00 |
|  | n_inact × sort_type | 18 | 570 | 3.16 | 0.00 |
| rHebb<br>(n = 20) | n_inact | 9 | 570 | 83.7 | 0.00 |
|  | sort_type | 2 | 570 | 4.24 | 0.01 |
|  | n_inact × sort_type | 18 | 570 | 0.98 | 0.48 |

**Figure 7:**
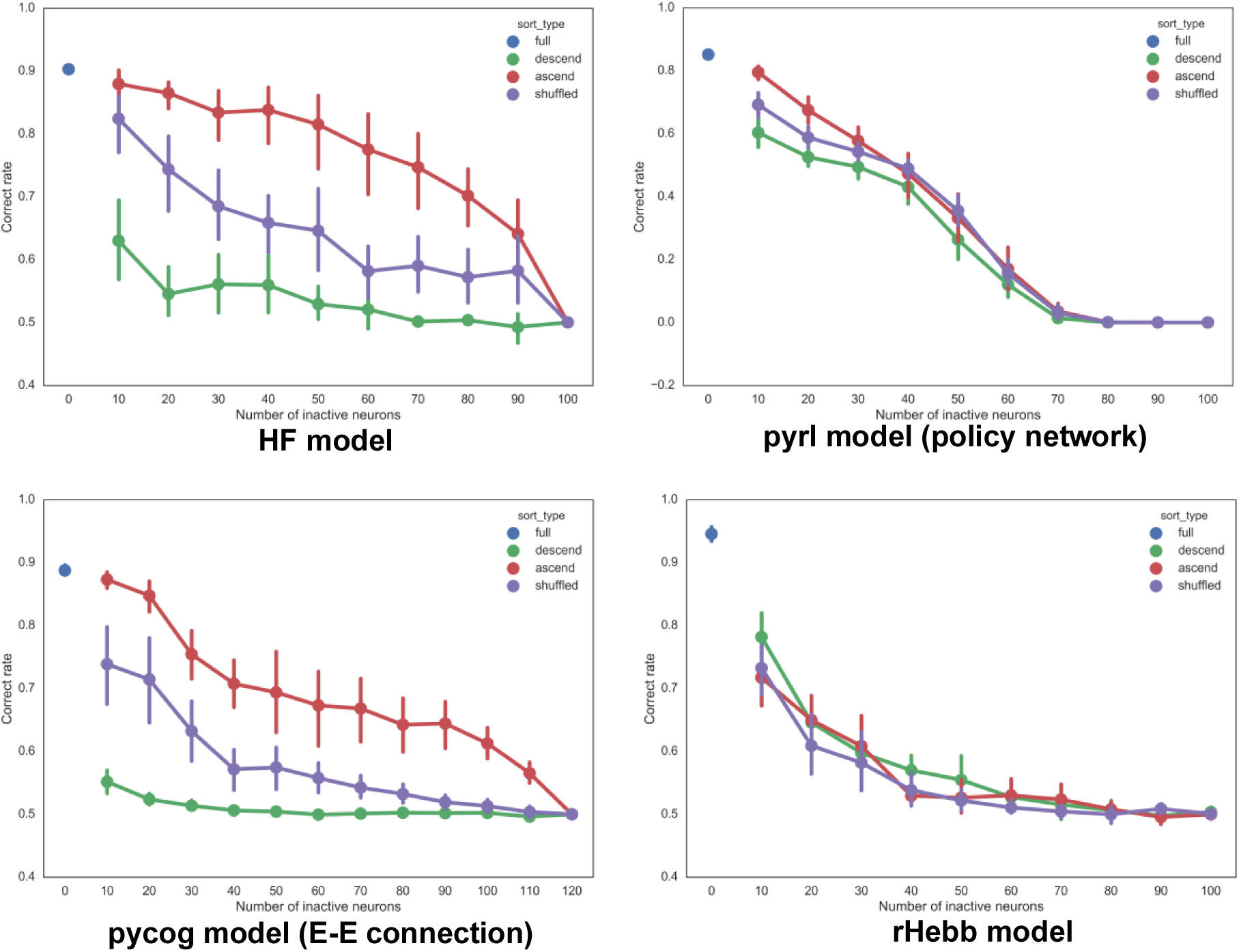
Accuracies of task performances in the unital inactivation experiment.

The features of weight change distribution could be altered depending on the initial network state or behavioral tasks. We checked the weight distributions for the different behavioral tasks and initial conditions. Most of them showed comparable results, which have highly skewed post-unital mean weight change distributions and high-kurtosis weight change distributions (Figure5-Figure supplement 2 and 3, Figure5-Table supplement 1 and 2). The weight distribution of the pycog model was initially a uniform distribution; however, after learning, it became a highly skewed distribution. In contrast, the weight distribution of the pyrl model initially had a Gaussian distribution, and its distribution remained Gaussian after learning (Figure5-Figure supplement 3). We also studied the rHebb model, which initially had a uniform distribution. However, the learning of the task could not be sufficiently converged (data not shown).

## Discussion

We analyzed static network structures of four RNN models for performing context-dependent integration tasks. We found that the structure of synaptic changes has some common features, i.e., high skewness in post-unital mean weight change distributions (Figure 5) and high kurtosis in weight change distributions (Figure 6), in contrast to no similarities in the structures learned last (Figures 3 and 4). The high-plasticity units were large contributors to the behavioral performance tasks in most models (Figure 7). These detailed data acquisitions and modifications may not have been achieved without the computational models. Additionally, as far as we know, detailed analyses of the RNN PFC models, such as the ones reported here have not been performed previously. These results indicate that plastic changes may be more important to perform cognitive tasks than accomplished network structures, which may have an impact on both computational and biological viewpoints of future research.

**Figure 3:**
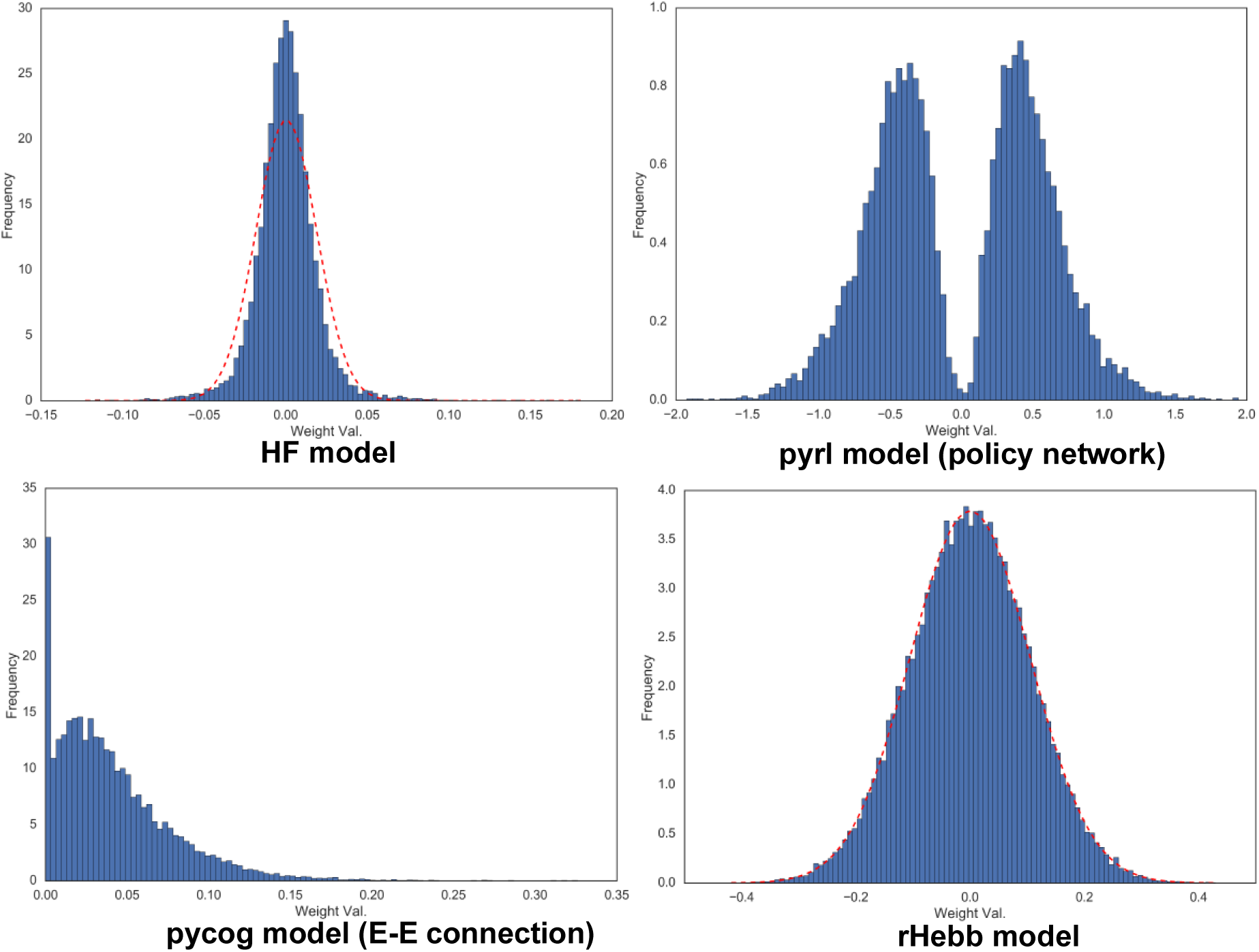
Weight distribution of each system after learning. The dotted red line in the HF and rHebb models represents a normal distribution with mean and sigma similar to the weight values

We found long-tailed distributions of plastic changes at the neuronal unit and synapse level as common features in all the models. In the biological brain, it is known that gene expression, which induces plastic changes in the neuron, is sparse in the cerebral cortex and hippocampus. In addition, it has been hypothesized that the plastic changes of a small neuronal population mainly represents learning and memory, namely, the engram hypothesis (Hebb, 1949; Tonegawa, Pignatelli, Roy, & Ryan, 2015). Additionally, at the synapse level, only a small population showed plastic changes with learning (Hayashi-Takagi et al., 2015; Yang, Pan, & Gan, 2009). In the neural network, protecting high-informative synapse while learning a new task can also prevent catastrophic forgetting of previously learned tasks (Kirkpatrick et al., 2017). Most of our results have comparable properties with the results of biological studies at both the neuronal (unital) and synaptic levels (Figure 5 and 6) and support the engram hypothesis (Figure 7).

Regarding the learned network structure, we could not find any similarities among the RNN models. It is well-known that in real brains, the distribution of excitatory postsynaptic potential is a log-normal distribution (reviewed in Buzsáki & Mizuseki, 2014). This corresponds to the weight distributions of the HF (Figure 3) and pycog models (Figure5-Figure supplement 3). Both models showed a long-tailed shape distribution regardless of the initial distribution setting. A previous study reported a long-tailed post-mean unital distribution of learned networks with the HF method (Chaisangmongkon, Swaminathan, Freedman, & Wang, 2017). Furthermore, excitatory neurons in the biological cortex form subclusters with mutual connections (reviewed in Harris & Mrsic-Flogel, 2013). Weight changes of pycog showed a high synaptic plasticity between high-plasticity units (Figure 5A), which could be the basis of the mechanism to construct the neural subclusters. Comparing these observations with the biological brain, HF and pycog models, followed by pyrl, were more similar to the animal brain, and rHebb was the most dissimilar. The result was opposite to our assumption that rHebb and pyrl models are more biologically plausible than pycog and HF models.

One potential reason why HF and pycog were found to be more similar to the real brain is the relationship between spike-timing-dependent plasticity (STDP) and SGD such that the dynamics of these two can be similar during learning (Bengio, Mesnard, Fischer, Zhang, & Wu, 2015; Xie & Seung, 2000). STDP is a major plastic feature in the biological brain (reviewed in Feldman et al., 2012) and it is thought to be one of the reasons for the log-normal synaptic distribution (Buzsáki & Mizuseki, 2014). If this is true, the pycog (adopting SGD) and HF models (HF, an advanced method of SGD) are more biologically plausible as synaptic update mechanisms, and it is reasonable to think that they are more similar to the biological brain than the other models in the last structure. A spiking RNN model adopting STDP can also perform similar cognitive tasks (Klampfl & Maass, 2013).

The pyrl model also adopted SGD, and the basic assumption of the eligibility trace in the rHebb model (*Equation 17* of this paper) seems to be similar to the assumption in the STDP model (Formula (1) or (2) in Bengio et al., 2015). However, the pyrl and rHebb models showed milder kurtosis in weight change distribution (Figure 6) and relative equal contribution of each unit to the task performance (Figure 7) compared with the HF and pycog models. The major difference between the two groups was whether the learning method was reward-based reinforcement or error (between the reference signal and system output)-based supervision. We thought that the difference in learning processes of each method, particularly during the initial stages, could cause the differences of the distributions. Error in supervised learning is the largest and relatively constant during the initial stages of learning. The responsible synapses for the error seem to be fixed. Conversely, rewards in reinforcement learning were occasionally supplied depending on the stochastic fluctuation of the network activities. The network updates based on stochastic activities and can randomly scatter contributions of the synapses to task performance, particularly during the initial learning stages. With these views, the network might get sparse by supervised learning (and STDP), while there may be less cell or synapse loss by reinforcement learning.

Another possible explanation for the highly skewed synaptic weight distribution observed for the biological brain is that it results from multiple experiences through life. Based on our results, the synaptic change distributions of all models were long-tailed distributions (Figure 6). Repeating the learning processes, even with mild plastic changes, such as those in the rHebb model, can result in a long-tailed distribution in the real brain.

Our results revealed that the features in plastic changes of weight values were more general than the learned weight structure itself. We can easily control weight distribution shapes of the learned structure of the computational models with additional terms of objective function such as the regularization term (Lee, Battle, Raina, & Ng, 2006), which can make the distribution sparse and long-tailed. These are possible reasons why the last structure learned is less common among the models than plastic changes, even though the initial distributions and terms are critical for the learning efficiency. The other properties of neural plasticity have been previously reported in the biological brain, such as synaptic homeostasis (reviewed in Pozo & Goda, 2010) and stabilization (reviewed in Koleske, 2013), which should correspond to regularization (Lee et al., 2006) and elastic weight consolidation (Kirkpatrick et al., 2017) in the neural network methods, respectively. Considering these factors, we can establish more biologically plausible computational models in future studies.

Common properties in synaptic change may be explained by the superlinearity of either the activation function or learning rule in each model. In the rHebb model, the superlinear functions of the learning rule (but not linear or sublinear ones) lead to sparse and precise synaptic change (Miconi, 2017), which can establish the high-kurtosis synaptic change distribution. Moreover, the pycog and pyrl models use the so-called rectifier activation function (ReLU) to calculate the firing rate. Such rectifier units act as a superlinear function and make neural activity sparse (Glorot, Bordes, & Bengio, 2011), resulting in only a few neurons to be active and plastic.

We focused on the context-dependent integration task to know the necessary structures involved in the process of achieving flexible cognition. Our findings, however, seem to apply more generally to the learning tasks than the cognitive flexibility task (Figure5-Figure supplement 2). In this study, we only analyzed the synaptic weight structures of RNN models. As a future work, we plan to also analyze the dynamics of unital activities while performing a task and the underlying learning process. These analyses will give us further insights on how the networks encode and establish the task information. Furthermore, theoretical investigation is also needed to understand the meaning of our findings and further develop the RNN models. Recent innovations in optimization RNN methods allowed the systems to perform cognitive tasks similar to human and model animals, which allows us to compare the processes of the biological and computational brains (Cadieu et al., 2014; Carnevale, de Lafuente, Romo, Barak, & Parga, 2015; Mante et al., 2013; Sussillo, Churchland, Kaufman, & Shenoy, 2015; Yamins et al., 2014). Merging such knowledge, methods, and ways of thinking in both fields that study cognitive tasks must improve our understanding of how our brain works.

## Materials and methods

### Model descriptions

The parameter settings were set to default values based on previous reports and scripts (Mante et al., 2013; Miconi, 2017; Song et al., 2016, 2017). All the models were constructed as follows:

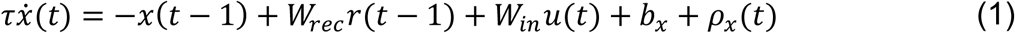

where τ ∈ ℝ is the time constant, 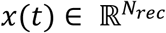 corresponds to themembrane potentials of recurrent neuronal units at discrete time step *t*, *N*_*rec*_ is the number of recurrent units, 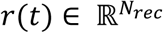represents the firing rate and is calculated by applying the rectified linear activation function (*r*(*t*) = *x*(*t*) for *x*(*t*) Ȁ and *r*(*t*) = 0 otherwise; for the pycog and pyrl models) or the hyperbolic tangent function (*r*(*t*) = tanh(*x*(*t*)); for the HF and rHebb models) to *x*(*t*). *u*(*t*) ∈ ℝ^*N*^*in* is an (external) task input consisting of sensory and contextual information, *N*_*in*_ is the number of input units (four channels in the HF and rHebb models and six channels in the pycog and pyrl models), *w*_*rec*_ ∈ 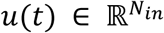 and *W*_*in*_ ∈ 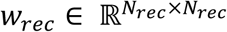 are the synaptic weight matrices from recurrent and task inputs to each recurrent unit, respectively. 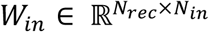 is the offset constant of recurrentunits, and 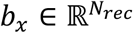 is the noisy fluctuation of each unit following a Gaussian distribution. For the pyrl model, we used a gated recurrent unit (Chung, Gulcehre, Cho, & Bengio, 2014) and *Equation 1* was modified (see section pyrl model). Note that we used *N*_*rec*_ =100 for HF, *N*_*rec*_ = 150 for pycog (120 excitatory and 30 inhibitory units), and *N*_*rec*_ = 200 for rHebb. For pyrl, we used *N*_*rec*_ = 100 for the policy network and *N*_*rec*_ = 100 for the baseline network.

The use readout units of HF, pycog, and pyrl were as follows:

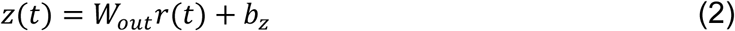

where 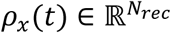 is the output of the system, *N*_*out*_ is the number of the output units (one channel in the HF model, two channels in the pycog model, and three channels in the pyrl model),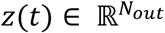 is the synaptic weight matrix from recurrent units to readout units, and 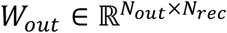 his the offset constant of the readout unit. By contrast, the rHebb model used an arbitrary recurrent unit as an output. The choices of the system were represented as the signs of the output unit (*z*(*t*) in the HF model and arbitrary unit of *r*(*t*) in the rHebb model; one channel in total) or the highest channel (among two channels in the pycog model and among three channels in the pyrl model) of the output units. Only the pyrl model had additional choices (like “stay”) to choice 1 and choice 2. The *N*_*in*_, *N*_*rec*_, and *N*_*out*_ of each model are summarized in Table 6.

**Table 6:** Number of input, recurrent, and output units of each model

| Model | $N_{in}$ | $N_{rec}$ | $N_{out}$ |
| --- | --- | --- | --- |
| HF | 4 | 100 | 1 |
| pycog | 6 | 150 (Ex: 120, Inh: 30) | 2 |
| pyrl (policy) | 6 | 100 | 3 |
| rHebb | 4 | 200 | 1 (an arbitrary unit) |

### Task descriptions

The task inputs, *u*(*t*) in *Equation 1*, consisted of two sets of sensory and two sets of contextual information. Sensory inputs were as follows:

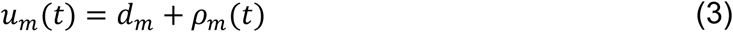

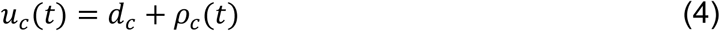

where *u*_*m*_(*t*) ∈ ℝ^1^ ^or^ ^2^ and *u*_*c*_(*t*) ∈ ℝ^1^ ^or^ ^2^ are the motion and color sensory inputs, respectively, *d*_*m*_ ∈ ℝ^1^ ^or^ ^2^ and *d*_*c*_ ∈ ℝ^1^ ^or^ ^2^ are the offsets, and *ρ*_*m*_(*t*) ∈ ℝ^1^ ^or^ ^2^ and *ρ*_*c*_(*t*) ∈ ℝ^1^ ^or^ ^2^ are Gaussian noises with zero mean. The amplitudesof *d*_*m*_ and *d*_*c*_ represent motion and color coherence. Choice 1 or choice 2 was represented as a plus or minus signs of *d*_*m*_ and *d*_*c*_, respectively, in the HF and rHebb models (two channels in total), and as independent channel inputs in the pycog and pyrl models (four channels in total). The contextual information was modeled with two inputs, *u*_*cm*_(*t*) ∈ {0,1} and *u*_*cc*_(*t*) ∈ {0,1}: *u*_*cm*_(*t*) = 1 and *u*_*cc*_(*t*) = 0 in the motion context and *u*_*cm*_(*t*) = 0 and *u*_*cc*_(*t*) = 1 in the color context at every time step *t*.

### HF model

The HF model was implemented based on a previous study (Mante et al., 2013). HF optimization was mounted with modifying scripts written by BL Nicolas on Github (https://github.com/boulanni/theano-hf) (Boulanger-Lewandowski, Bengio, & Vincent, 2012).

HF optimization (Martens, 2010; Martens & Sutskever, 2011; Shewchuk, 1994) was processed by minimizing the following objective function *ε*(θ).

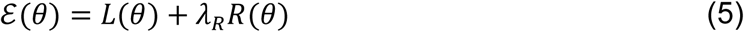

where

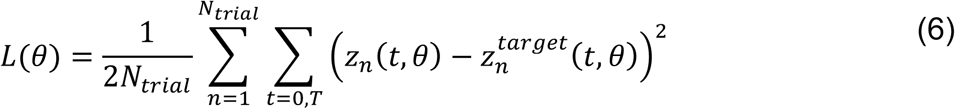

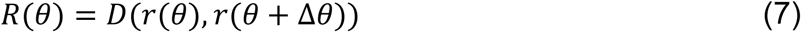

Note that *θ* is the vectorized parameter for the optimization, including*W*_*rec*_, *W*_*in*_, *W*_*out*_, *b*_*x*_ and *b*_*z*_, where *L*(*θ*) indicates the error between the target 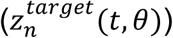 and actual output (*zn*(*t*, *θ*))of the system at the first and last timepoints (*t* = 0, *T*) of all trials (*N*_*trial*_). *R*(*θ*) is the term of structural damping to prevent disruptive changes of recurrent unital activities because a small perturbation of the recurrent units can result in a very large output difference (Martens & Sutskever, 2011). λ_*R*_ > 0 determines the degree of the *R*(*θ*) penalty, and its value is determined using the Levenberg-Marquardt algorithm (Nocedal & Wright, 1999), and *D*(*r*(*θ*), *r*(*θ* + ∆*θ*)) is the distance (cross entropy in our script) between outputs of recurrent units with parameter *θ* and *θ* + ∆*θ*. The target outputs 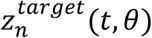 of the last time points (t = *T*) were considered correct by setting the target output 1 and −1 to correspond with choice 1 and choice 2, respectively, the target outputs and 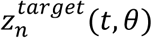 of the first time points (t = 0) were set to 0 regardless of the correct choices to fix the initial points of the output transients. The minimum of the object function was calculated with HFoptimization (Martens, 2010; Martens & Sutskever, 2011).

The values *N*_*in*_ = 4, *N*_*rec*_ = 100, and *N*_*out*_ = 1 were used in the HF model. Because of memory capacity, the time step was 25 times larger than the original setting (Δt = 25 ms). The standard deviations of noises reported previously (Mante et al., 2013) were decreased in the present study to the following equation: Std[*ρ*_*x*_] = 0.004 in *Equation 1* and Std[*ρ*_*m*_], Std[*ρ*_*c*_] = 0.04 in *Equation 3* and *Equation 4*, respectively.

### Pycog model

The pycog model was obtained from Github (https://github.com/xjwanglab/pycog) and was run in its original setting (Song et al., 2016). This system learns tasks with modified SGD (Pascanu et al., 2013) by minimizing the following objective function *ε*(*θ*).

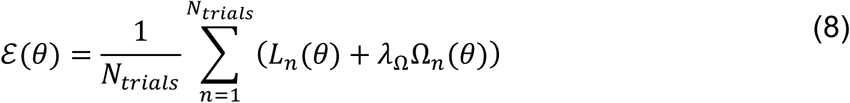

where *θ* is the vectorized parameter set for the optimization, *N*_*trials*_ is thenumber of trials, and *Ln*(*θ*) is the error between the actual and target outputs(*z*_s_ (*t*, *θ*) ∈ ℝ^2^ and 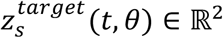, respectively) through trial (*T*) and the number of output units (*N*_*out*_) given by

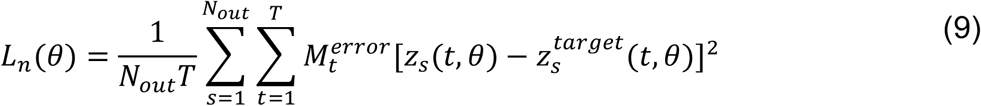

where 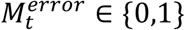 is the error mask consisting of 0 or 1 and determines whether the error at time point *t* should be taken into account (in a context-dependent integration task; only the last output is considered). Ω*n*(*θ*) in *Equation 8* is a regularization term for preserving the size of the gradients as an error during learning processes, and λΩ determines the effects of the regularization.

The values *N*_*in*_ = 6, *N*_*rec*_ = 150 and *N*_*out*_ = 2 were used in the pycog model. Of note, this system includes both excitatory and inhibitory units at an excitatory to inhibitory ratio of 4:1, indicating that the number of excitatory units is 120 and that of inhibitory units is 30. For our network analyses, we mainly used E-E connections. The initial weight distribution of the default setting is a gamma distribution and multiplied signs depending on the input unit types (excitatory or inhibitory). Applying a uniform distribution as an initial weight distribution, the minimum and maximum are 0 and 1, respectively.

### pyrl model

The pyrl model was also obtained from Github (https://github.com/xjwanglab/pyrl) and was run in its original setting (Song et al.,2017). The network consists of policy and baseline RNN, in which the nodes are gated recurrent units (Chung et al., 2014). We modified *Equation 1* as follows:

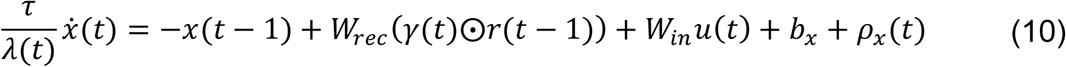

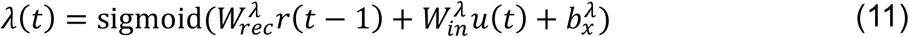

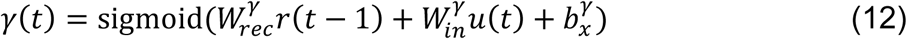

Where 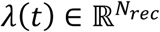 and 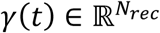 are the updat and reset gates, respectively, 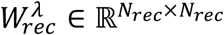 and 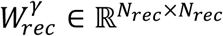 are the weight matrices from recurrent units,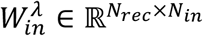 and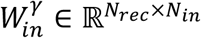 are the weight matrices from task inputs, 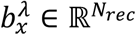 and 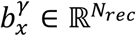 are the offset constants of update gates λ(*t*) and *γ*(*t*), respectively, and ⊙denotes an element-wise product (Hadamard product). λ(*t*) and *γ*(*t*) are nonlinearized with sigmoid function. A policy recurrent network takes sensory, context, and inner recurrent inputs and selects the next action at every time step. The outputs are normalized with the softmax function 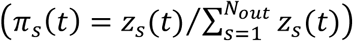. A baseline network receives recurrent unit activities (*r*(*t*)) and choice outputs (*π*(*t*)) of the policy network, and predicts the sum of reward values through the trial.

The policy network aims to maximize the expected future rewards, which areoptimized using the REINFORCE algorithm (Wierstra et al., 2010; Williams, 1992).

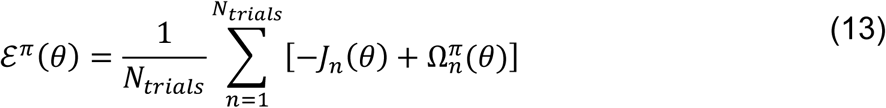

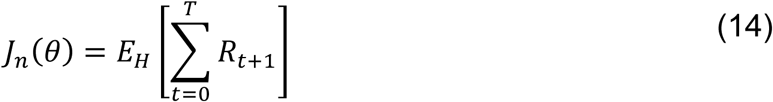

where *θ* is the vectorized parameter set of the policy network for the optimization, *ε^π^*(*θ*) is the objective function of the policy network, *N*_*trials*_ is the number of trials, *j*_*n*_(*θ*) is the expected reward prediction, 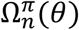 is regularization term, and *E*_*H*_ represents the expectation of reward *R*_*t*_ over all possible trial histories *H*.

The baseline network minimizes the difference between the actual and the estimated reward values through the trial.

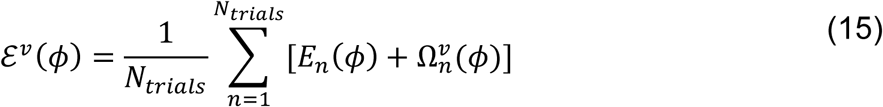

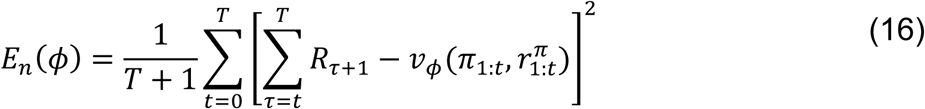

where *φ* is the vectorized parameter set of the baseline network for the optimization, *ℰ*^*ν*^(*θ*) is the objective function of the baseline network, *E*_*n*(_*θ*) is the error between the actual reward *R*_τ_ (a correct decision is rewarded with *R*_τ_ = 1, if incorrect *R*_τ_ = 0, and the duration of breaking the fixation before the decision is negatively rewarded with *R*_τ_ = −1), 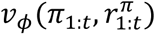 is the expected (readout) reward prediction of the baseline network under recurrent unit activities 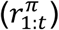 and choice (*π*_1:t_) of the policy network through a trial (*T*), and 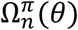 is regularization term of the baseline network. This system uses Adam SGD (Kingma & Ba, 2015) with gradient clipping (Pascanu et al., 2013) for optimization.

The values *N*_*in*_ = 6 (task inputs), *N*_*rec*_ = 100, and *N*_*out*_ = 3 (choice) were used in the policy network, and the values *N*_*in*_ = 103 (*r*(*t*) and *π*(*t*) of the policy network), *N*_*rec*_ = 100, and *N*_*out*_ = 1 (readout reward prediction) were used in the baseline network. We used the policy network for the main analysis of this study because the baseline network was not involved in performing the task (the baseline network is critical for the optimization of the system). The default initial weight distribution was obtained from the gamma distribution with multiplying signs randomly in both policy and baseline networks. The plastic synapses were set at 10% of all synapses and the other synaptic weights were fixed in default through the training. We only used plastic synapses for the weight change distribution analysis. This system had three output choices (choice 1, choice 2, or stay) although the other models had only two choices (choice 1 or choice 2). When a normal distribution was used as an initial weight distribution, the mean and standard deviation are 0 and 1, respectively

### rHebb model

TherHebbmodelwasobtainedfromgithub(https://github.com/ThomasMiconi/BiologicallyPlausibleLearningRNN) and wasbasically run in its original setting (Miconi, 2017). The network pools Hebbian-like activity in every time step as follows:

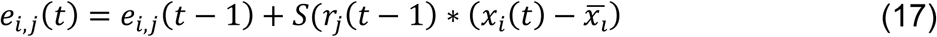

where *e*_*i*,*j*_(*t*) is the accumulated eligibility trace of the synapse *i* (pre) and *j* (post), *S* is the monotonic superliner function (in this case *S* = *x*^3^), *r*_*j*_(*t*) is the output of unit *j*, *x*_*i*_(*t*) is the membrane potential of unit at *t*, and 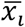 is theshort-time running average of *x*_*i*_. The synaptic weights are modulated with the pooled value and reward error at the end of every trial as follows:

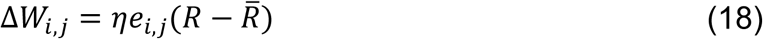

where ∆*W*_*i*,*j*_ is the change of synaptic weight between *i* and *j, η* is the learning rate, *R* is the reward from the environment (absolute difference from optimal response is supplied as negative reward), and 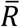 is the average of previously received reward. Three units are used as constant input unit in default. The output and constant input units are excluded from the weight change distribution analysis.

The values *N*_*in*_ = 4 and *N*_*rec*_ = 200 and *N*_*out*_ = 1 were used in rHebb model. The output of this system is an activity of an arbitrarily chosen unit from the recurrent units.

### Statistical analysis of distributions

Python libraries, numpy, scipy, statsmodels, matplotlib, seaborn, and Jupyter were used for statistical analysis. Shapiro-Wilk normality test was applied to check the normality of the distributions, which was implemented as a scipy function. Kurtosis and skewness were tested as scipy functions, stats.kurtosistest and stats.skewtest (https://docs.scipy.org). One-way ANOVA, two-way ANOVA and multiple comparisons (Tukey honestly significant difference) were performed with the python library, statsmodels (http://www.statsmodels.org)

### Neuronal unit inactivation

Scripts were modified as shown below. Selected unit outputs were set to 0 (pycog, pyrl, and rHebb model) or a constant value (HF model, *b*_*x*_) in every recurrent loop. The number of inactivation units was increased in increments of 10 and the orders were sorted in ascending, descending, or shuffled order of the postsynaptic mean of absolute synaptic weight changes (Figure 5A). The trial settings, including the number of trials and offset and noise settings of sensory inputs, also followed default conditions in each script.

### Source scripts

Codes used in this work are available at https://github.com/sakuroki/flexible_RNN.

## Acknowledgment

We thank Naoki Hiratani for technical advice on RNN optimization, Taro Toyoizumi for valuable contributions to the discussion, and Shigeyoshi Itohara for allowing us to publish this study based on work done at his laboratory.

### Author contributions

SK, Conceptualization, Software, and Writing; TI, Theoretical analysis and Writing

## Supplemental Figures

**Figure3-Figure supplement 1:**
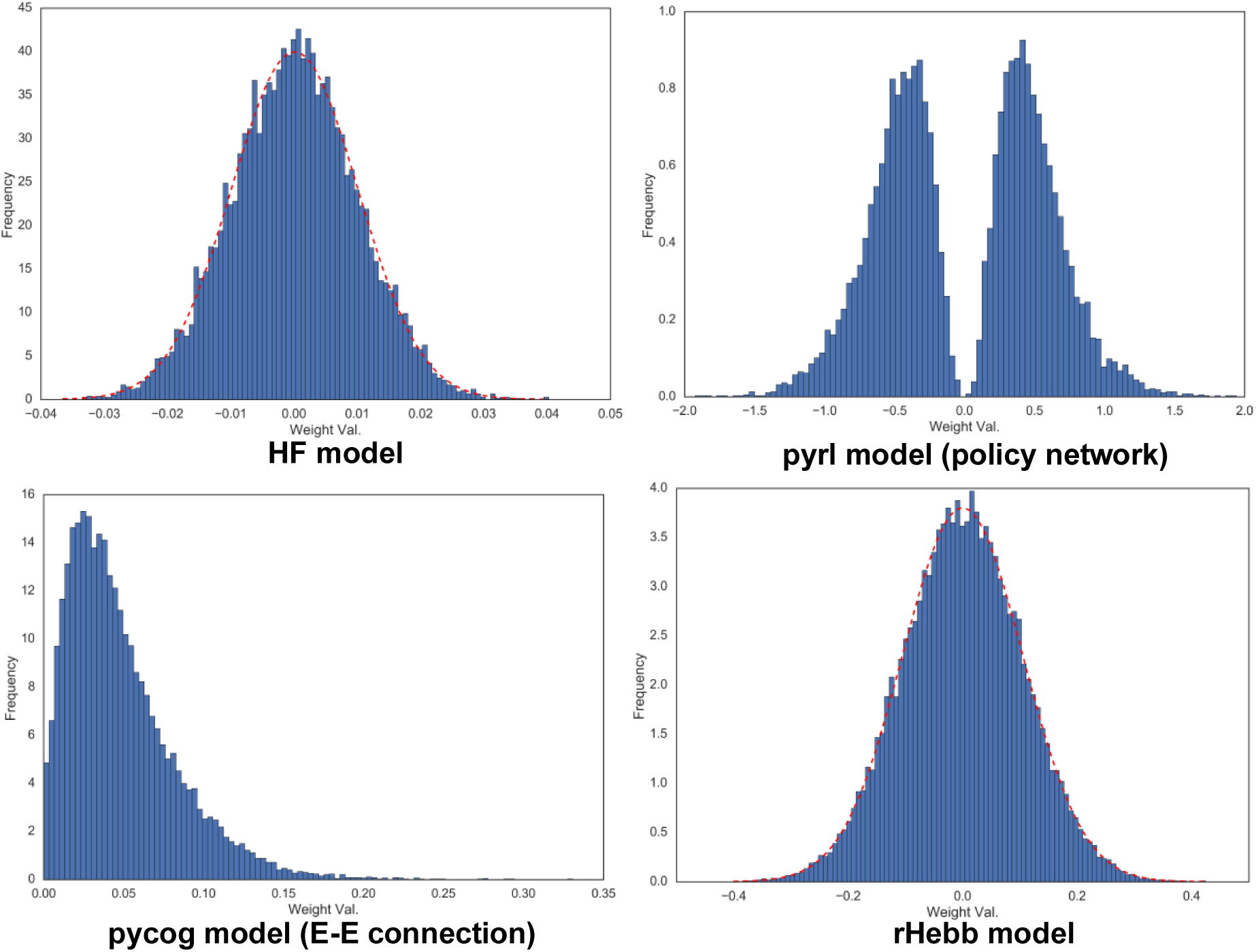
Initial synaptic weight distribution of each model.

**Figure5-Figure supplement 1:**
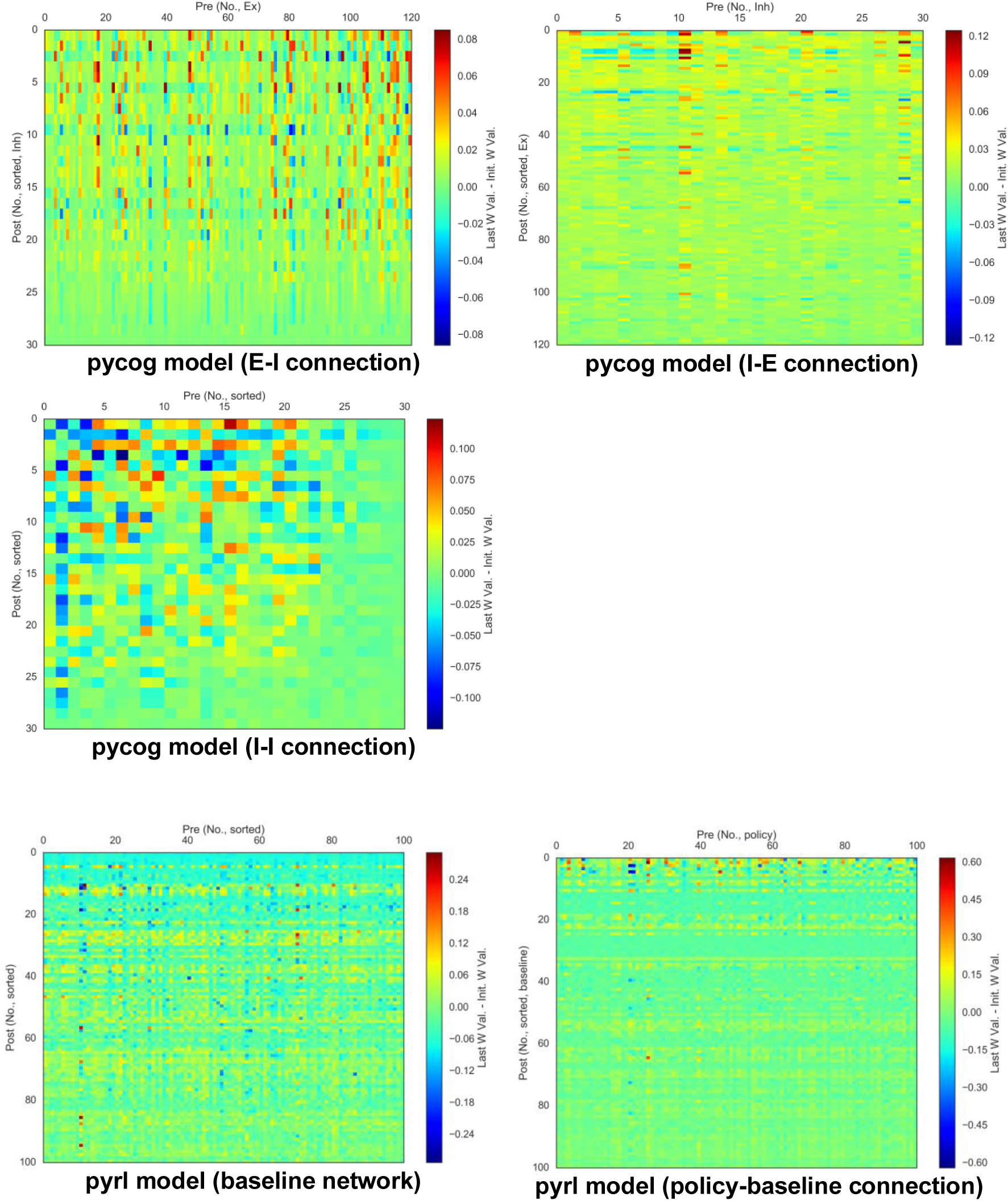
Weight change plots of pycog I-I, I-E, E-I, pyrl baseline-baseline and policy-baseline connections, sorted by post-unital mean weight changes.

**Figure5-Figure supplement 2:**
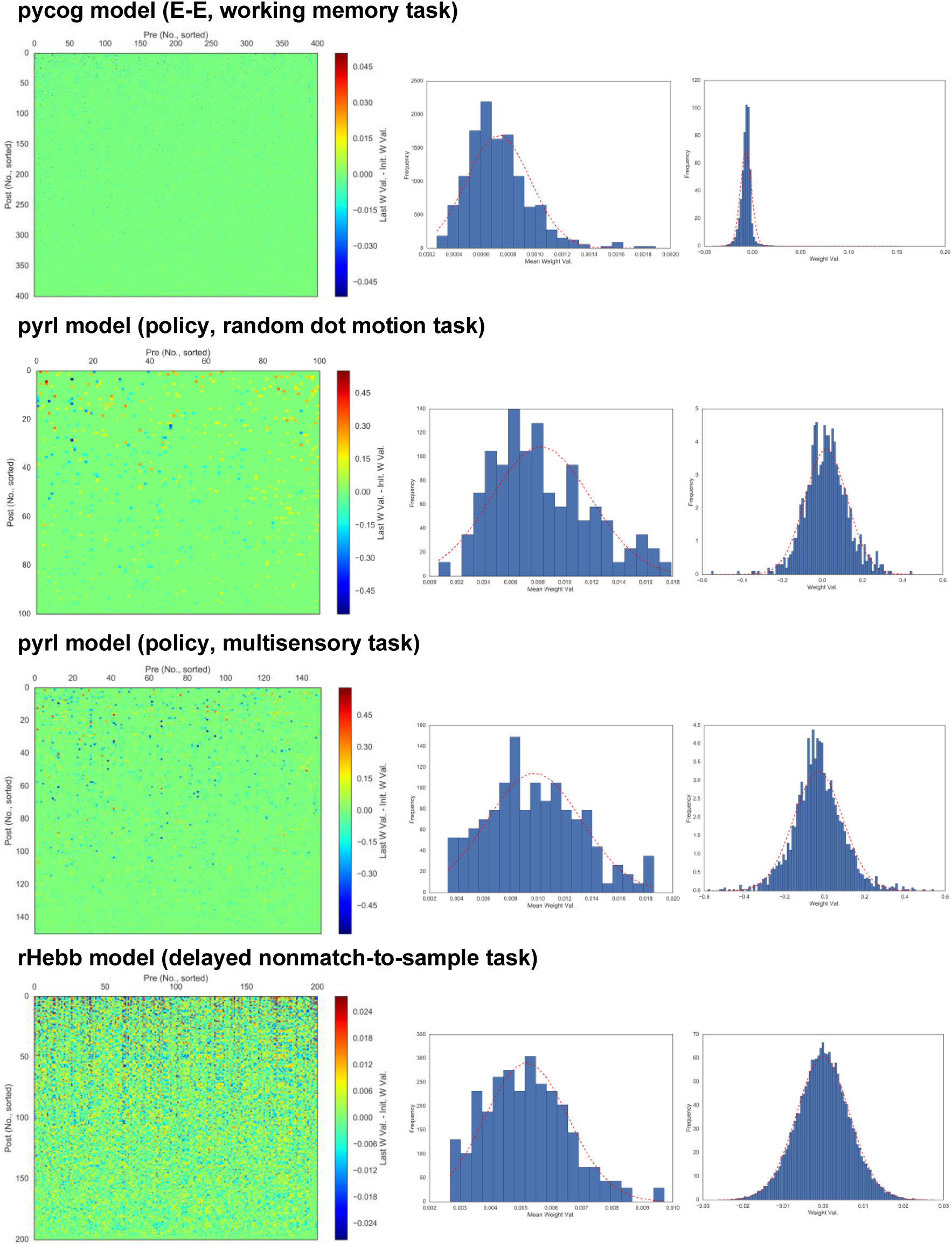
Sorted weight changes plot (left), post-unital mean weight change (middle) and weight change (right) distributions performing different cognitive tasks. References for each task are shown below (working memory (Romo, Brody, Hernández, & Lemus, 1999), random dot motion (Gold & Shadlen, 2007), multisensory (Raposo, Kaufman, & Churchland, 2014), delayed nonmatch to sample (Simola et al., 2010)).

**Figure5-Figure supplement 3:**
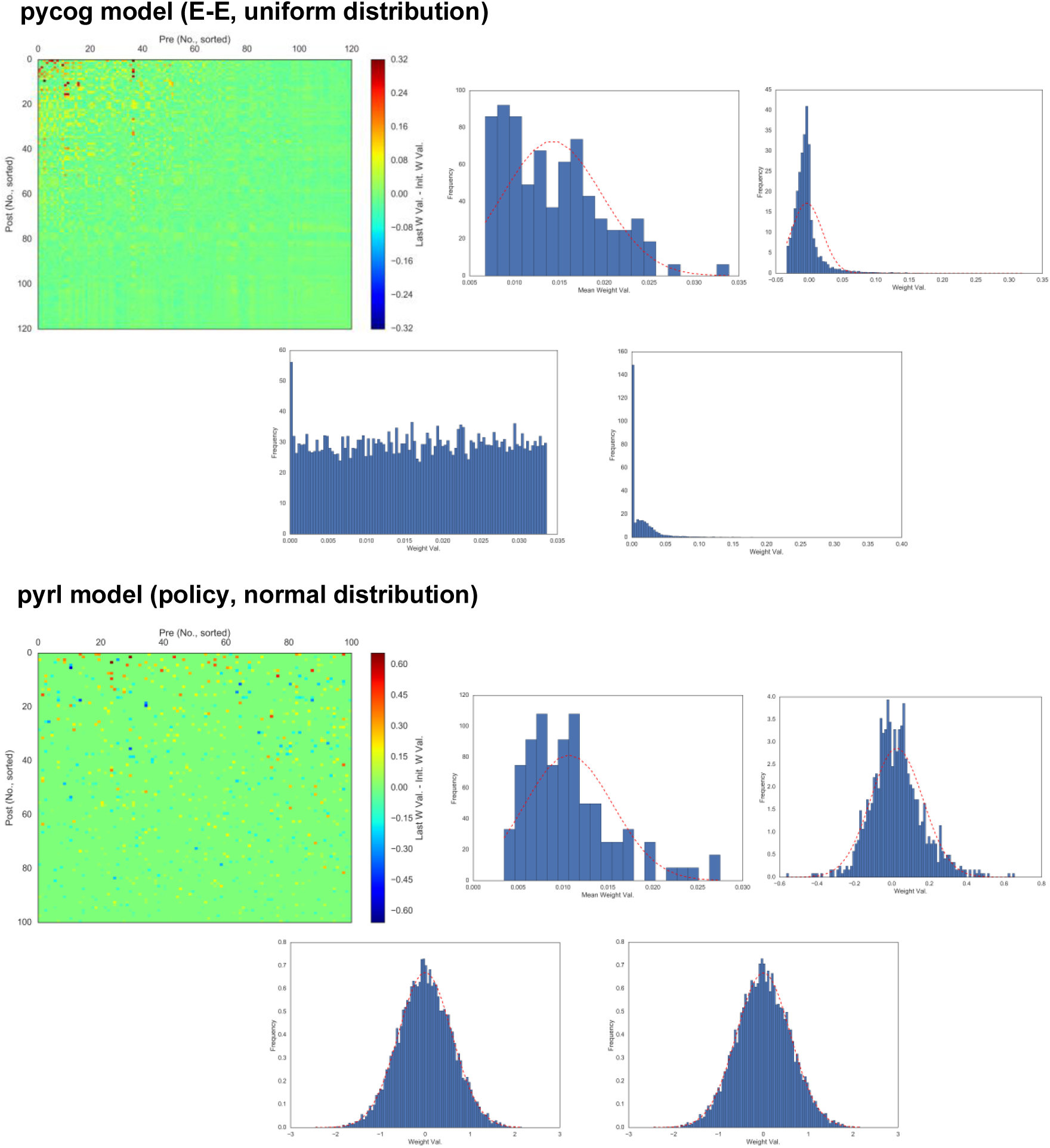
Sorted weight change plot (upper left), post-unital mean weight change (upper middle), weight change (upper right) distributions with different initial weight distributions. Lower left and right panels show the initial and last weight value distributions, respectively.

## Supplemental Tables

**Figure5-Table supplement 1:**
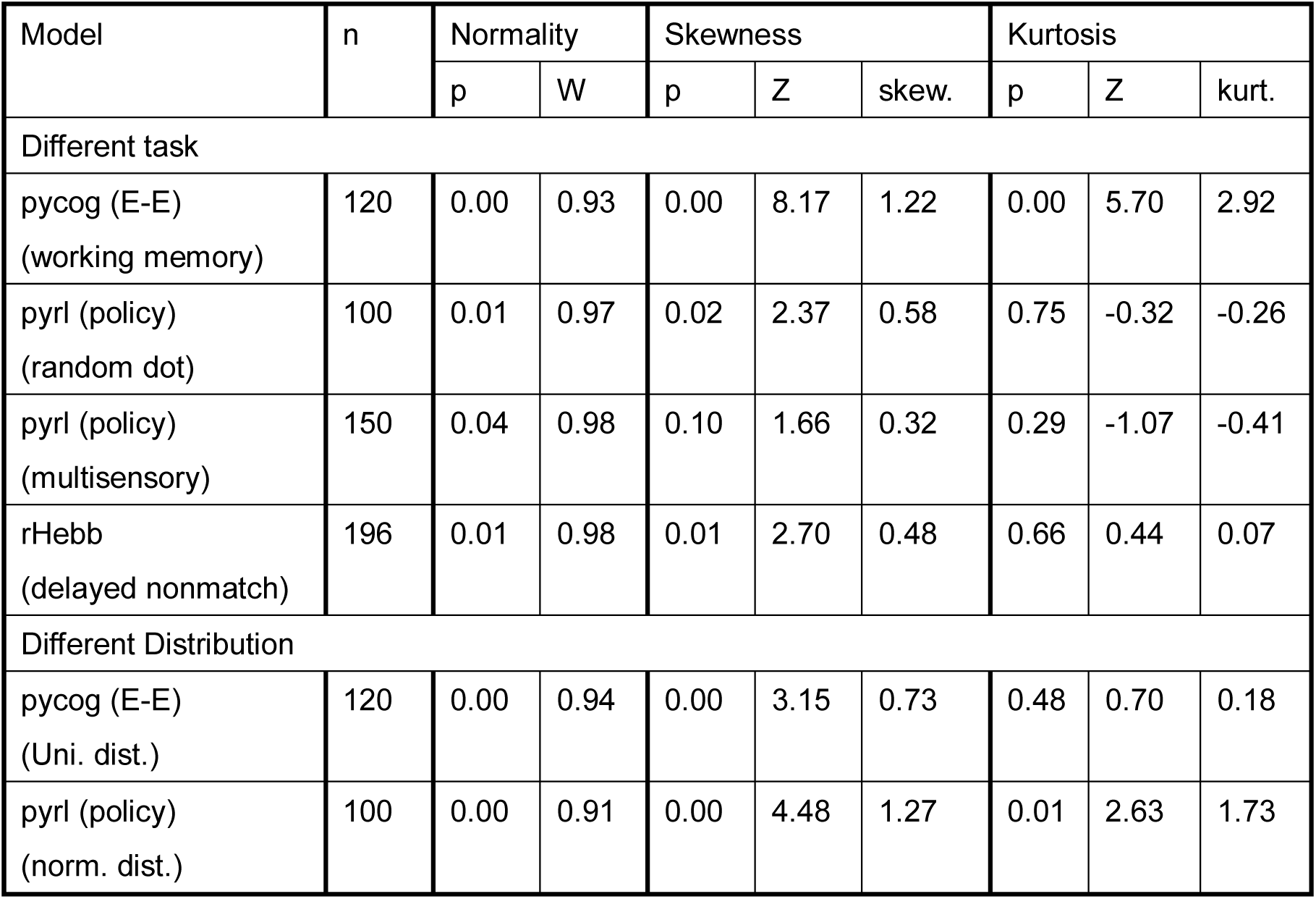
Properties in post-unital mean weight change distributions in different initial states and tasks

**Figure5-Table supplement 2:**
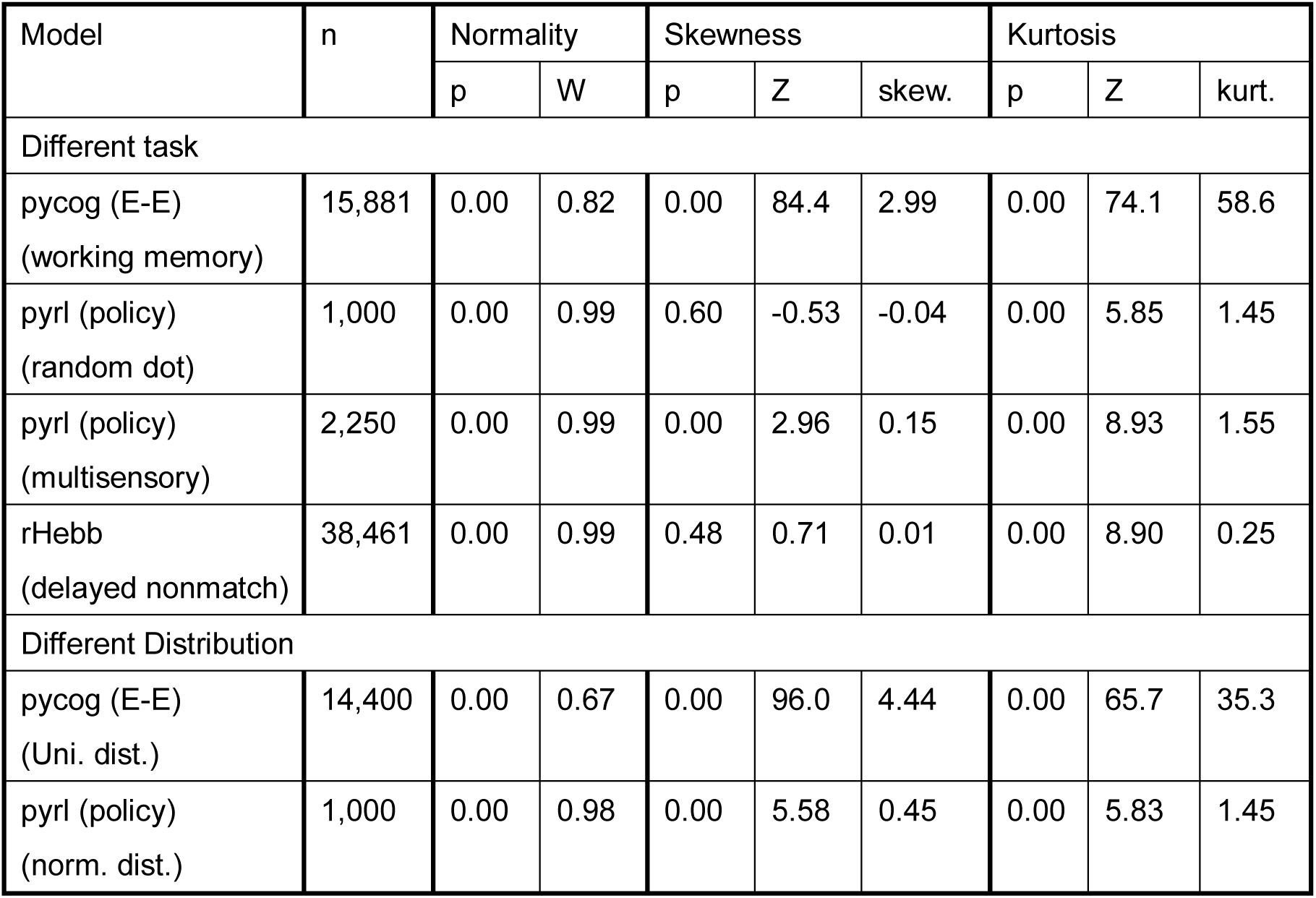
Distribution properties in weight changes in different initial states and tasks

